# The physiological response in the mammary gland to *Escherichia coli* infection reflects a conflict between glucose need for milk production and immune response

**DOI:** 10.1101/603845

**Authors:** Shlomo E. Blum, Dan E. Heller, Shamay Jacoby, Oleg Krifucks, Uzi Merin, Nissim Silanikove, Gabriel Leitner

## Abstract

Upon entry into the mammary gland, mammary pathogenic strains of *Escherichia coli* rapidly grow using milk as a nutrient source, in a manner highly dependent on the expression of the ferric-dicitrate system by these bacteria. Intra-mammary challenge with distinct mammary pathogenic *E. coli* (MPEC) strains result in development of clinical acute mastitis, with peak bacterial counts in milk at 16-24 h post-challenged and profound immune changes found in the milk. The main biochemical changes measured in milk were partially in accord with lipopolysaccharide-induced mastitis, with increased glucose-6-phosphate and lactate dehydrogenase activity or prolonged lactate dehydrogenase, and Glu6P/Glu alterations. Changes also reflect physiological responses to inflammation in the mammary gland, as in the balance between aerobic and anaerobic metabolism (citrate to lactate ratios). Some alterations measured in milk resolved with days after challenge but other remained for above one month, regardless of bacterial clearance. The results suggest that *E. coli* mastitits can be divided into two stages: an acute, clinical phase, as an immediate response to bacterial infection in the mammary gland and a chronic phase, independent of bacteria clearance, and in response to the tissue damage caused in the first phase.

## Introduction

*Escherichia coli* is the main pathogen causing acute mastitis in cattle, a disease of significant economic and animal welfare impact in dairy production worldwide. Upon entry into the mammary gland, mammary pathogenic strains of *E. coli* rapidly grow using milk as a nutrient source, in a manner highly dependent on the expression of the ferric-dicitrate system by these bacteria [1-3]. The resulting infection leads to an acute inflammation that causes a sharp decrease in milk yield and deep changes of milk properties and constituents [4]. Even after clinical healing and infection clearance in the mammary gland indicated by cessation of bacterial secretion in the milk, milk yield and quality might remain impaired for a long period in a significant number of affected cows. The degree and length of the post-acute inflammation effect on udder health and milk quality vary greatly and may depend on the health status of the individual animal, time of diagnosis and treatment, as well as the infecting *E. coli* strain [5].

Recently, intra-mammary challenge with distinct mammary pathogenic *E. coli* (MPEC) strains (VL2874, VL2732 and P4) and with the non-mammary pathogenic *E. coli* strain K1, was conducted in order to study if the dynamics and intensity of the immune response in bovine mammary glands is different, depending on *E. coli* strains [4]. Cellular and chemokine responses and bacterial culture follow-up were performed for 35 days post-challenge. Cows challenged by any of the MPEC strains developed clinical acute mastitis with peak bacterial counts in milk at 16-24 h post-challenge. During the 35 days of the study, differences were found in the intensity and duration of the response in various of the parameters studied, namely the somatic cells count (SCC), leukocytes distribution, secreted cytokines (TNF-α, IL-6 and IL-17) and levels of membrane TLR4 on leukocytes in milk.

Although a lot has been studied about the immune and inflammatory reaction to *E. coli* in the mammary gland, less is known about the physiological and biochemical changes that occur in the gland and in the milk during and following intra-mammary infections by these bacteria. In a previous study [6], it was shown that under acute mammary inflammation induced by intra-mammary challenge of cows with *E. coli* lipopolysaccharide (LPS), the passage of glucose-derived carbons shifts to the pentose phosphate pathway, leading to a cell metabolism shift to glycolysis at the expense of mitochondrial respiratory activity. A negative correlation between milk secretion and lactose, glucose and citrate concentrations and inflammation parameters was described. In parallel, the concentrations of glucose-6-phosphate, malate, oxaloacetate, lactate and pyruvate, and the activities of the respective enzymes glucose-6-phosphate dehydrogenase, malate dehydrogenase and lactate dehydrogenase increased and, in general, positively correlated with inflammation parameters. A main difference between LPS and live bacteria intra-mammary challenge is the fact that bacteria represent a continuous stimulant of the mammary immune system due to bacteria replication in the gland, which may remain for days after the infection commence, whereas a single LPS injection would be a one-time stimulus. There is limited knowledge on the physiological and biochemical changes induced by live MPEC and how these changes truly correlate to LPS-induced mastitis. The objective of the present work was to study the physiological and biochemical responses in milk of cows challenged with three distinct MPEC strains, as presented by Blum et al. [4]. The basis for the use of three different MPEC strains was to represent *E. coli* involvement in distinct presentations of bovine mastitis.

## Materials and Methods

### Animals

Ten Israeli Holstein cows at the Agricultural Research Organization (A.R.O.) dairy farm at lactations 1-6, 133-442 days in lactation, and with milk yield of 22-38.4 L/day were used for intra-mammary challenge with mammary pathogenic strains of *E. coli*. The cows were milked thrice daily (05:00, 13:00 and 20:00) in a dairy parlor equipped with an on-line computerized AfiFarm Herd Management data acquisition system, including the AfiLab milk analyzer, which provides on-line data on gross milk composition (fat, protein and lactose) and milk conductivity as measure of mastitis (Afimilk, Afikim, Israel; http://www.afimilk.com). The average milk yield in the herd was >11,000 L during 305 days of lactation. Animals were fed a typical Israeli total mixed ration containing 65% concentrate and 35% forage (17% protein). Food was offered *ad lib* in mangers located in sheds. At the beginning of the study, all glands of the 10 cows were free of infection as tested by three consecutive bacteriological tests and had SCC < 1×10^5^ cell/mL milk. Animal experimentation was approved by the Committee of Animal Experimentation of the A.R.O., the Volcani Center, Bet Dagan and followed the committee ethical guidelines (Permit no. 59315).

### Bacteria challenge

Three different *E. coli* strains were used for the intra-mammary challenge: VL2874 (isolated from per-acute mastitis), VL2732 (isolated from persistent mastitis) and P4 (widely used as model strain for mammary pathogenic *E coli*). Strains typing, phenotypic and genomic characteristics were described before [2, 7, 8]. Bacteria were recovered from frozen stock (kept at −80 °C in a mixture of brain-heart infusion and 25% glycerol) on blood agar (Tryptose Blood Agar Base; Becton-Dickinson, Sparks, MD, USA, with 5% washed sheep erythrocytes) and incubated aerobically at 37 °C overnight. For inoculum preparation, bacteria were harvested and washed in pyrogenic-free saline (PFS), suspended in PFS and stored at 4 °C for 10 h. Aliquots were taken for colony forming units (CFU) count on blood agar. CFU number was adjusted with PFS right before challenge, aiming to an inoculum of ~ 50 CFU in 3 mL. Final CFU number in the inoculum ranged from 10-30 CFU/gland, as determined from aliquots separated just prior to challenge. The small CFU counts in the inoculum aimed to better represent a natural infection in which small numbers of bacteria are expected to entr the mammary gland and initiate the infection.

### Study layout

All cows were challenged in one gland while all the other three glands served as a control. Before challenge, milk yield, milk composition, SCC and leukocytes distribution were determined at −2 days and on day 0, separately for the intendent challenged gland and composite milk of the other three glands. At the day of challenge, after morning milking, a single gland (left right in nine cows and front right in one cow) was infused via the teat canal with one of the *E. coli* strains (3-4 cows each). Clinical symptoms were assessed for the first 24 h, including measurements of rectal temperature every 4 h. After challenge, milk yield was recorded separately for the challenged gland and the three control glands. Milk was sampled at: 4, 8, 12, and 16 h post-challenge, and then at 1, 2, 4, 7, 10, 14, 17, 21, 29, and 35 days post-challenge. A week after the termination of the experiment (at day 45 post-challenge) the cows were slaughtered in an abattoir. The challenged and control teats and up to ~10 cm of parenchyma above were taken on ice to the laboratory for histological and PCR analyses for the presence of the bacteria.

### Histology

Samples for histological analysis were fixed in neutral buffered 4% formaldehyde and embedded in paraffin. Sections were cut (4 μm) and stained with hematoxylin and eosin (H&E). Slides were viewed using a light microscope (Nikon Eclipse E600, Melville, NY, USA), and images were captured using the MagnaFire Digital Camera (Optronics, Goleta, CA, USA) controlled by NIS-Elements F3.0 software (Nikon, Japan).

### Milk sample collection and analyses

For bacteriological tests, teats were cleaned, disinfected, foremilk was dispensed, and 3 mL samples of milk were aseptically collected in sterile assay tubes. For the other tests, the infected gland and then composite milk of the other three glands were milked separately into containers, milk volume was recorded and gently mixed and 0.5-1.0 L was taken for analysis as follows: SCC with the Fossomatic 360 (Foss Electric, Hillerød, Denmark) and gross milk composition, i.e., fat, protein, casein and lactose contents with the Milkoscan FT6000 (Foss Electric). These analyses were performed at the Israel Cattle Breeders Association Laboratory (Caesarea, Israel). Rennet clotting time (RCT) and curd firmness (CF) were tested using the Optigraph^©^ (Ysebaert, Frepillon, France) as previously described [9]. Fromase 15 TL (Gist-Brocades nv, Delft, The Netherlands) was diluted 1:100 from the stock solution and 0.5 mL/well were added to achieve clotting at about 900 s in non-infected milk. Tests were performed within 24 h after sample collection with milk stored at 4 °C. All the reagents for the below described procedures were purchased from Sigma (Rehovot, Israel). The concentrations of glucose (Glu), glucose-6-phosphate (Glu6P), Oxalaic acid (OA), malate (Ma), lactate (La), Pyruvate (Pyr) and citrate (Cit), were determined in skimmed milk and other food samples [10-12] by fluorometric methods in which the last stage was linked to formation of fluorochromophore as follow:

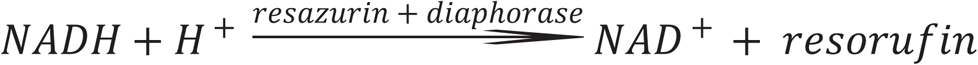

These fluorimetric procedures are insensitive to the disturbing effects of fat and casein micelles and the coupling to the formation of fluorochromophore increased their sensitivity. The limit of detection of these procedures was 10-50 µM and their sensitivity, residual standard deviation, was ± 2-5 µM.

The activity of malate dehydrogenase (MDH) and lactate dehydrogenase (LDH) in milk samples was done as follows. The final activity of diaphorase and the concentration of resazurin used for these enzyme-activity determinations were the same as used for the metabolite determinations. The basic reaction conditions for MDH activity in milk were as described by Silanikove et al. [6]. The differences in NADH concentration (between 10 and 1000 μM) at the linear stage of the reaction were divided by time and were converted to activity where 1 U of MDH will convert 1.0 μmol/L OA and NADH to L-malate and NAD^+^ per minute at pH 7.5 at 25 °C. The reaction conditions for LDH activity were as described by Larsen and Moyes [13]. One international unit of LDH activity was defined as the amount of enzyme that catalyzes the conversion of Pyr into La to generate 1.0 μmol/L of NAD^+^ per minute at 37 °C.

Nitrite concentration was determined by the fluorometric assay with DAN reagent as described by Sonoda et al. [14] after precipitating the casein by ultracentrifugation [15]. Lactic peroxidase (LPO) activity was assayed by the oxidation of 2,2 V-azino-bis (3-ethylbenzothiazoline-6-sulforic acid diammonium salt, as described by Silanikove et al. [15]. Catalase activity was determined with the aid of fluorescent substrate as previously described [16]. Additional sub-sets of milk samples were defatted under cold conditions and analyzed according to previously described procedures for albumin, lactoferrin and immunoglobulin type G (IgG) by ELISA assays [17-19].

### Statistical analysis

Ten dairy cows were tested for 38 different variables during the time course of the experiment. Each cow was tested in 20 different time points, from day 0 to 35 days post-challenge (DPC) during the study. The data sets of this study were analyzed using the PROC MIXED procedure of SAS (version 9.2, SAS Institute Inc., Cary, NC). A different dependent variable (38 variables) were tested over time with the general form: dependent variable = time cow(time). Consequently, in the description of the results, we focused on the presentation of the effects of time during the experiment on the level of a specific variable. Correlation models (Pearson correlation matrix) between all tested variables were performed using SAS Proc Corr. Data and are presented as means ± SEM.

## Results

### Clinical symptoms, bacteriology, SCC and milk yield

Rectal temperatures increased after 8 h, peaked at 12-16 h (39.2-40.1 °C) and decreased to normal after 16-24 h. Local clinical swelling was observed only in the challenged glands from 12-16 h and it persisted for 1 d (1 cow), 2 d (4 cows), 7 d (1 cow) and 35 d (4 cows). Each gland was inoculated with 10-30 CFU. *E. coli* bacteria were detected in milk as early as 4-8 h post-challenge and peak bacterial counts were observed after 16-24 h, reaching on the average up to ~ 1×10^7^ CFU per mL milk. Bacteria clearance was observed at 7 DPC, but in 50% of the glands bacteria were isolated from milk intermittently until the end of the study. Bacterial isolates from all cows at different time points were genotypically identical to the inoculated strains as tested by ERIC-PCR. Bacterial DNA was not detected in samples without bacterial growth in culture.

SCC in the infected glands increased significantly (*P*<0.001) within ~12 h days post-challenge (DPC) to over log 6 (1×10^6^ cell/mL). SCC peaked on day 1 to log 7 and remained at that level in all challenged glands up to 7 d, after which it started to decrease to ~log 6 until the end of the study (35 d) with no differences if bacteria was isolated or not (Fig. 1). The increase of SCC consisted mainly of leukocytes (CD18^+^ marked cells), mostly polymorphonuclear (PMNs) [4].

**Figure 1.**
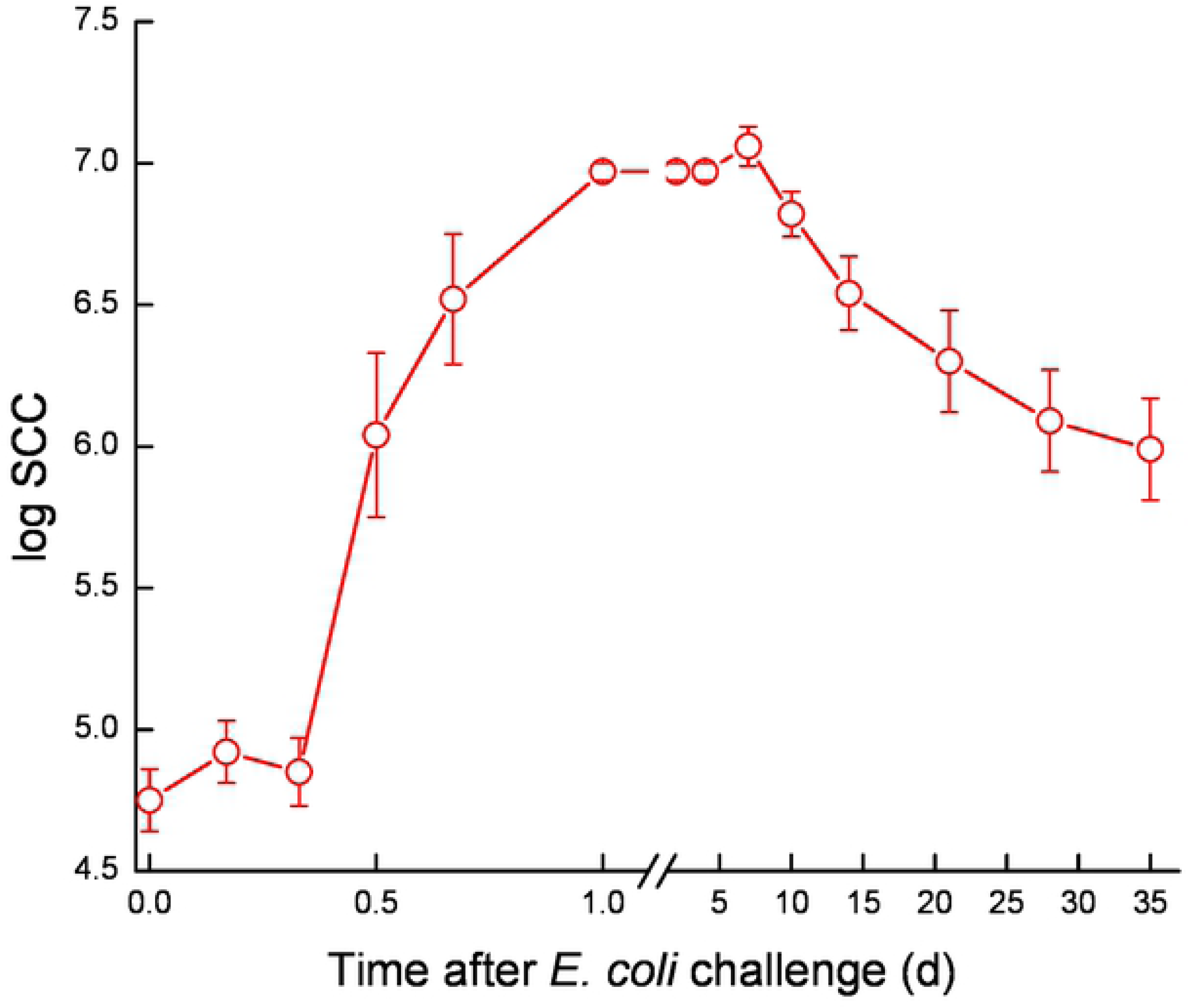
Mean and SE of somatic cell count in milk of the infected glands of 10 cows challenged with different mammary pathogenic strains of *E. coli*.

Milk yield was calculated as percentage of yield on day 0 (22-38.4 L/d; 100%). Milk yield of challenged glands was compared to composite milk of the control glands of each cow and to the total cow’s milk (Fig. 2). During 1-3 DPC, milk yield dropped to 10-25% in the infected glands and to 30-50% in the control glands. None of the challenged glands fully recovered during the study period (35 d), showing approximately 40-60% loss of milk yield, compared to ~20% loss in some control glands, while in others a compensation (gain in milk yield) was recorded. Overall, cow’s milk yield decreased by 10-30% at the end of the study (Fig. 2). The loss of milk yield per average cow during the 35 days of the study was 336 L (32%). Milk yield of infected glands significantly correlated with most of the parameters tested and significantly negatively correlated with SCC at the cow level (S1 Fig.).

**Figure 2.**
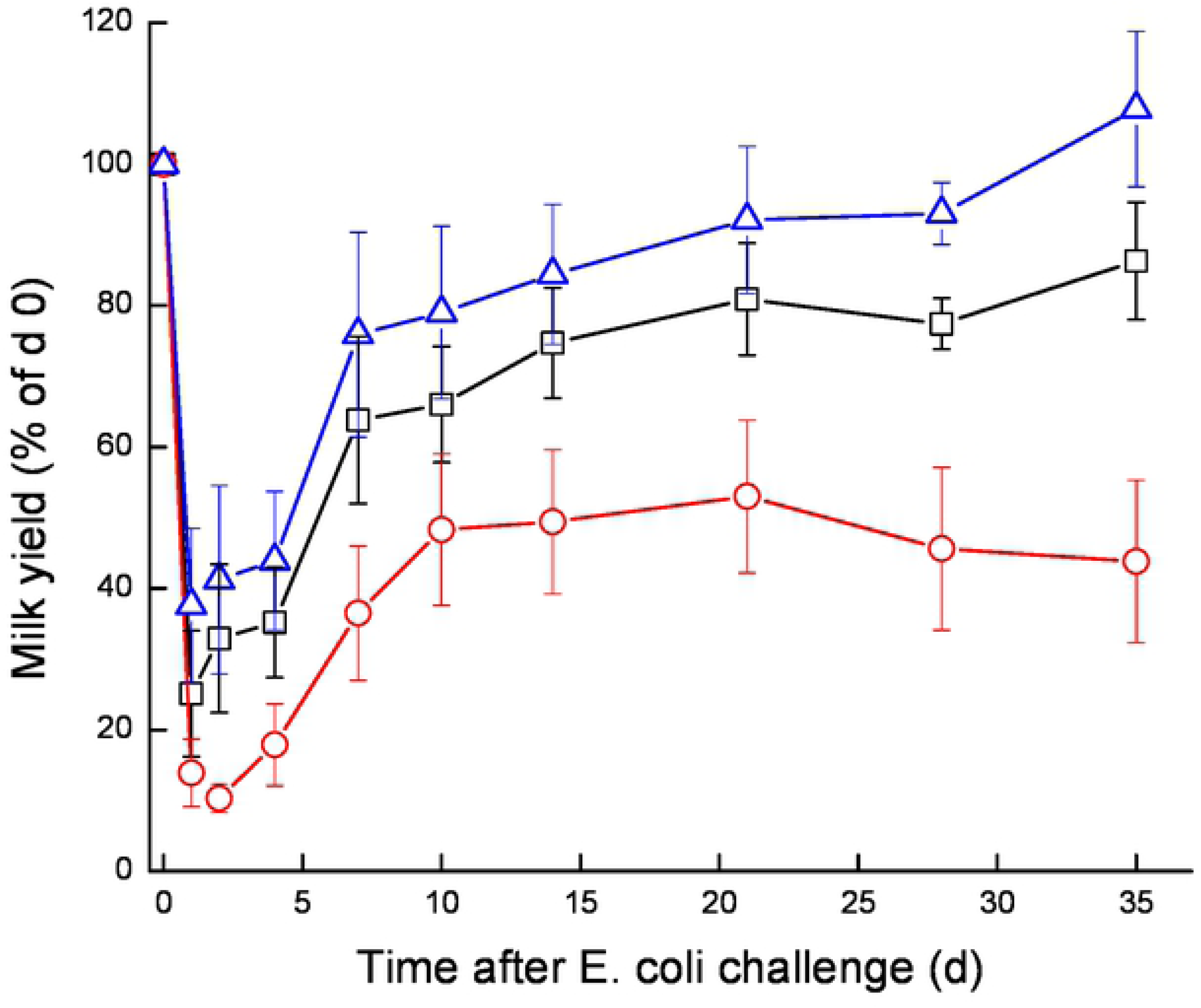
Mean and SE of milk yield measured separately for the infected glands (○), commingled milk of the 3 uninfected control glands (△) and the total cow’s milk yield (□) of 10 cows challenged with different mammary pathogenic strains of *E. coli*. Milk yield was calculated as percentage of day 0 (100%).

### Biochemical response to *E. coli* challenge

No significant results were found between the distinct MPEC strains; therefore, the following results are presented for the three MPEC strains combined.

The statistical significance (*P* values) of the ANOVA results for the Day effect, R^2^ and % variance between cows from the overall variance in the trial is summarized in Table 1. Fat in milk decreased during the first 2 h and then significantly increased (*P*<0.001) up to 7-10 DPC and gradually returned to pre-challenged level only at 35 DPC (Fig. 3). A significant decrease (*P*<0.001) in total protein was already noticed in 6-8 h DPC, after which total protein levels in milk increased up to a peak at 7-10 DPC and remained high till the end of the study, returning to pre-challenged levels at 35 DPC. In parallel, % casein decreased significantly from ~78% to ~68% between 8-24 h post-challenge, then gradually increased afterwards but remained significantly lower than pre-challenged levels until the end of the study (35d) (Fig. 3, Y2).

**Figure 3.**
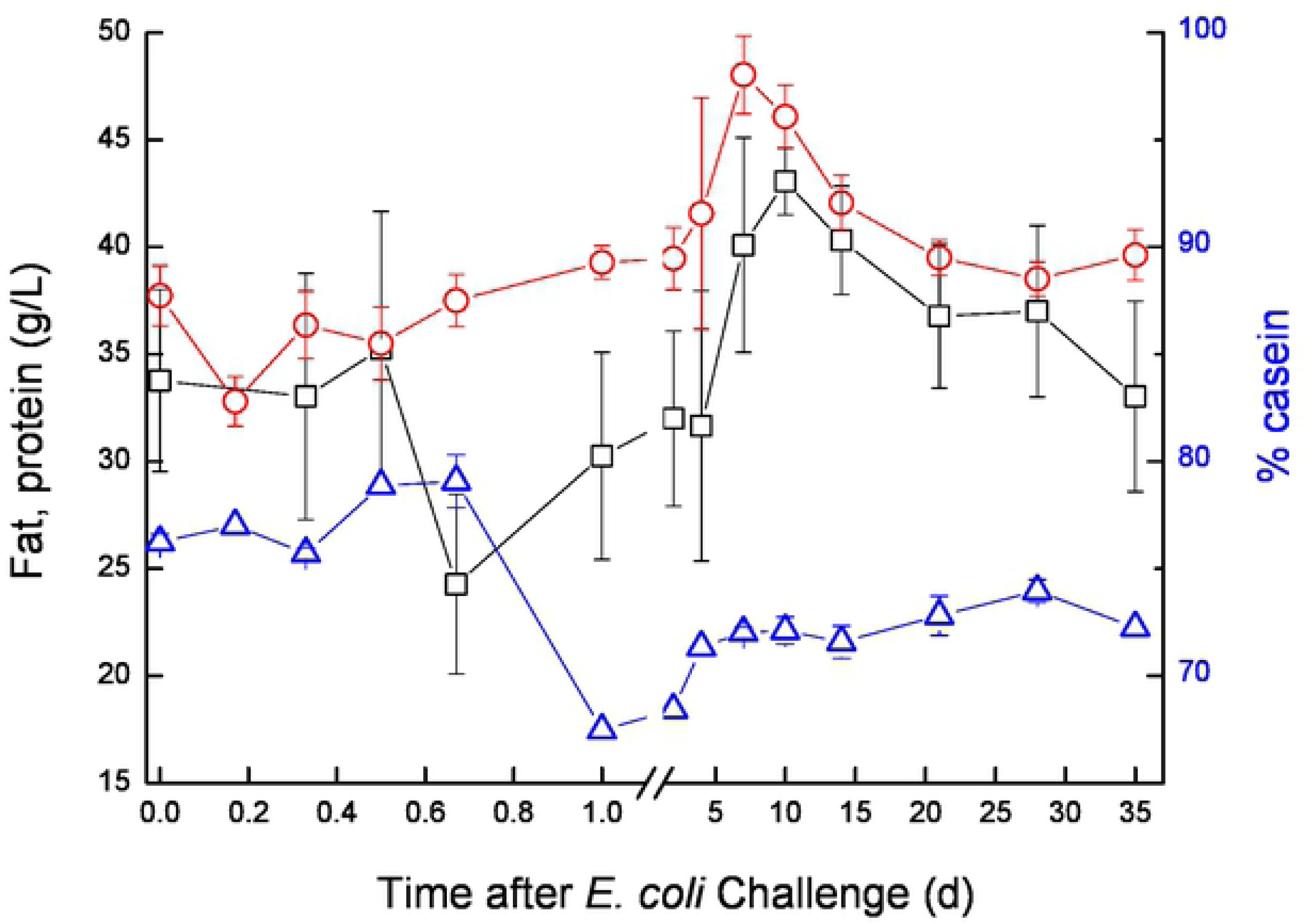
Mean and SE of fat (○; g/L), protein (□; g/L) and % casein (△), measured in the infected glands of 10 cows challenged with different mammary pathogenic strains of *E. coli*.

**Table 1.** The significance (*P* values) of the ANOVA results for the Day effect and the R^2^.

| Variable | R-Square (day) | P[*F*] | Variance between cows |
| --- | --- | --- | --- |
| Log SCC | 0.749 | <0.001 | NS |
| Milk (infected gland) L/d | 0.403 | <0.001 | NS |
| Milk (cow) L/d | 0.121 | NS | 15.2% |
| Fat (g/L) | 0.170 | NS | 18.8% |
| Protein (g/L) | 0.509 | <0.001 | 29.3% |
| % Casein | 0.399 | 0.008 | 76.1% |
| Lactose (g/L) | 0.521 | <0.001 | 25.8% |
| RCT (sec) | 0.755 | <0.001 | NS |
| CF (V) | 0.630 | <0.001 | NS |
| IgG (mg/mL) | 0.335 | <0.001 | NS |
| BSA (μM/mL) | 0.499 | <0.001 | NS |
| BLF (μM/mL) | 0.551 | <0.001 | NS |
| Lactate (μM) | 0.588 | <0.001 | NS |
| Malate (μM) | 0.589 | <0.001 | NS |
| Nitrite (nM) | 0.528 | <0.001 | NS |
| LDH (U/mL) | 0.648 | <0.001 | NS |
| Glucose (μM) | 0.228 | 0.016 | NS |
| Glu6P (μM) | 0.392 | <0.001 | NS |
| Glu6PD (μM/mL) | 0.362 | <0.001 | NS |
| Oxalate (mM) | 0.215 | 0.06 | NS |
| Lactoperoxydase (U/mL) | 0.227 | 0.04 | NS |
| Catalase (U/mL) | 0.267 | 0.007 | NS |
| Pyruvate (μM) | 0.337 | 0.001 | NS |
| Citrate (mM) | 0.318 | 0.004 | NS |

NS = Variance between cows was not significant ( $P > 0.05$ )

Milk coagulation properties assessed before challenge showed a mean RCT ~28 min and CF ~9 V. After challenge, milk coagulations properties were deeply affected and for the first 14 DPC, none of the milk of the challenged glands coagulated. From day 21, milk from 4-5 challenged glands coagulated but formed a weak coagulum, while the other cow’s infected glands milk did not coagulate until day 35 (Fig. 4).

**Figure 4.**
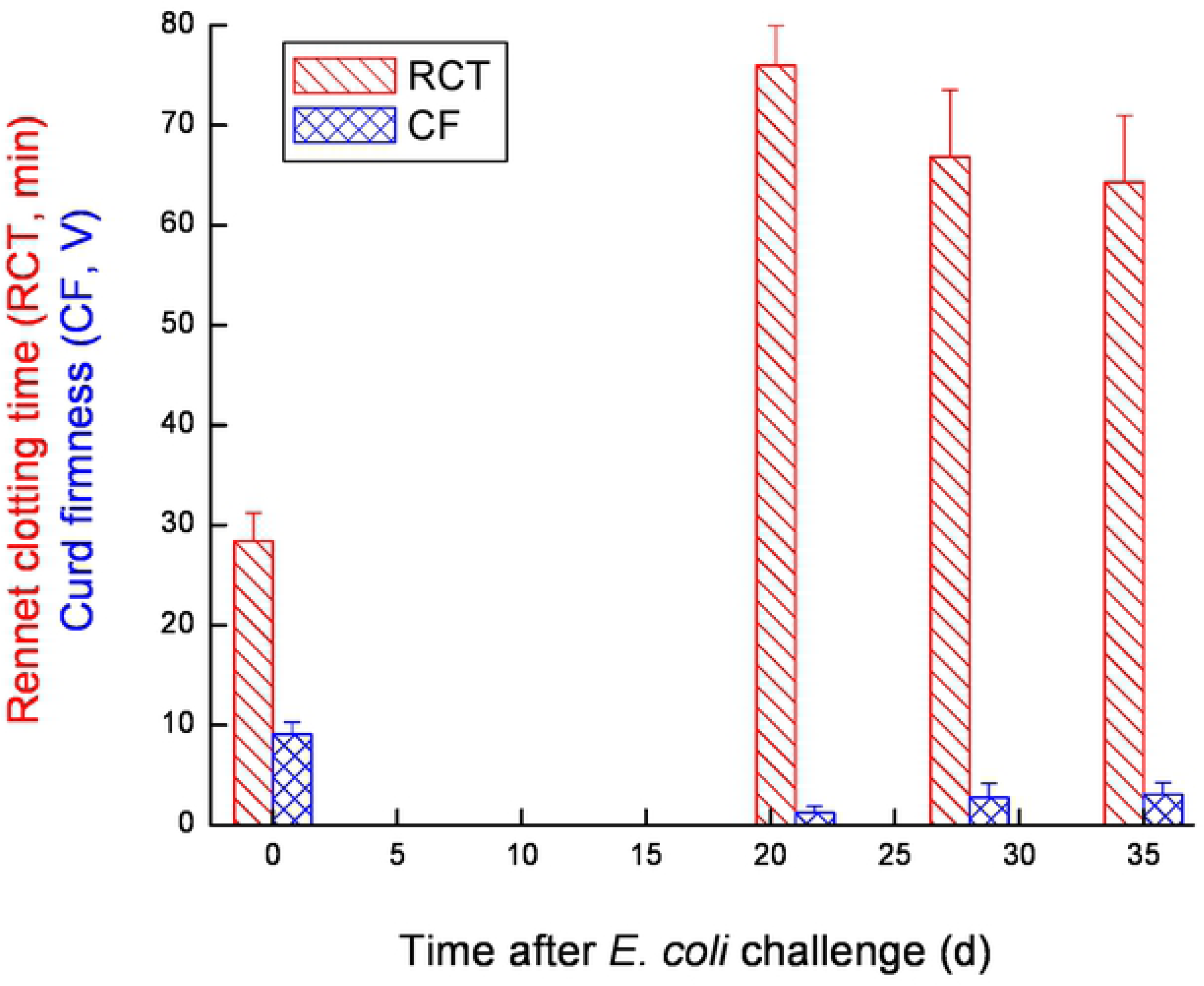
Mean and SE of Rennet clotting time (RCT) and Curd firmness (CF) measured in the infected glands of 10 cows challenged with different mammary pathogenic strains of *E. coli*.

Lactose in milk of the infected glands decreased at 12-16 h from ~50 g/L before *E. coli* inoculation to ~31 g/L and increased gradually level on days 28-35, but only for 60% of the infected glands, while it remained lower in the others (Fig. 5). The kinetics of changes in lactose concentration closely followed the kinetics of milk yield and the inflammatory response, as reflected by significant negative correlations with log SCC and the above-described inflammatory indices, and oxidative-stress markers (S1 Fig.). A significant positive correlation was found between protein level and inflammation parameters, while a significant negative correlation was found between % casein and lactose and inflammation (S1 Fig.). Regarding clotting parameters, a significant negative correlation was found between CF to inflammation and RCT, and significant positive correlation was found between CF to lactose and % casein (S1 Fig.), meaning that these parameters were similarly affected by the inflammation process.

**Figure 5.**
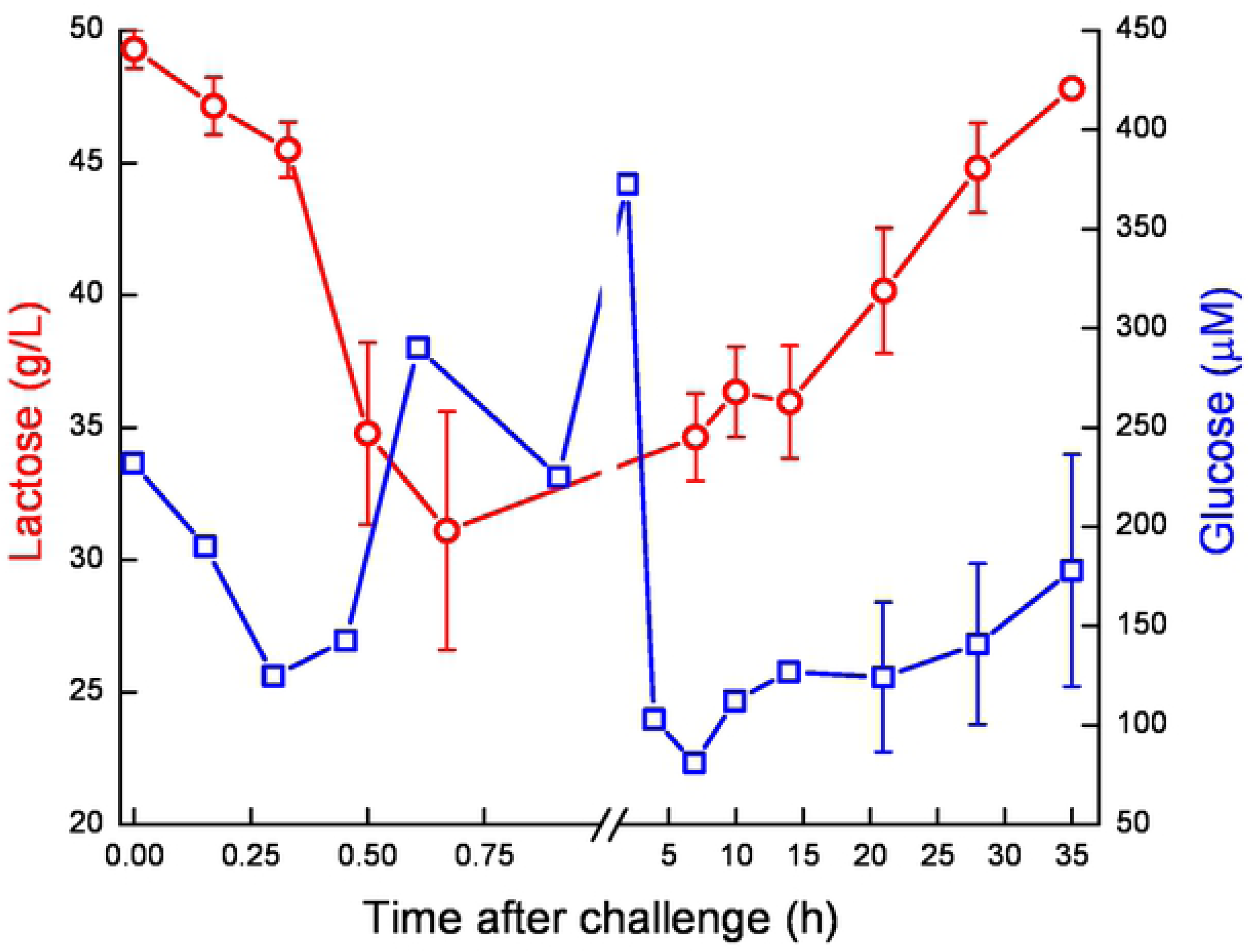
Mean and SE of lactose (○; g/L) and glucose (□; μM) measured in the infected glands of 10 cows challenged with different mammary pathogenic strains of *E. coli*.

The pre-challenged Glu concentration was ~289 μM. It increased on day 2 to ~370 μM followed by a sharp decrease on day 4 and gradually increase thereafter (Fig. 5). Apart from these changes, milk Glu concentration exhibited kinetics that closely followed that of the lactose. Glu6P in milk of the infected glands decreased at first on 12-16 h from ~49 uM before *E. coli* inoculation to ~33 μM, increased up to day 4. A sharp increase of Glu6P was observed on 4 DPC, coincidentally to a peak of Glu6P dehydrogenase activity (Glu6PD), decrease of Glu and subsequent peak of Glu6P/Glu ratio at day 4. This peak decreased on 10 DPC and Glu6PD activity gradually returned to pre-challenged levels, as well as Glu6P and the Glu6P/Glu ratio, which then fluctuated until the end of the study (Fig. 6A, B).

**Figure 6A-B.**
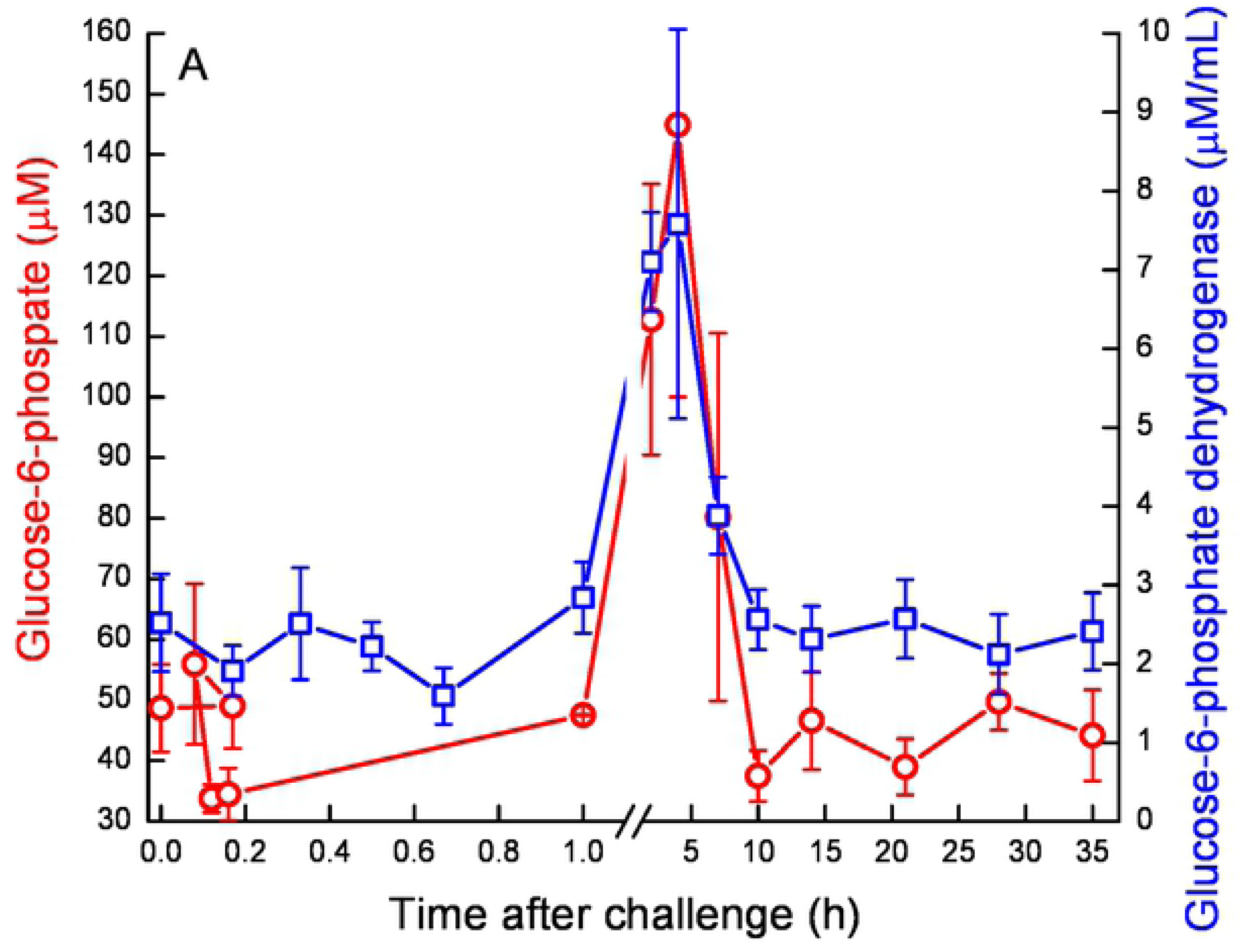
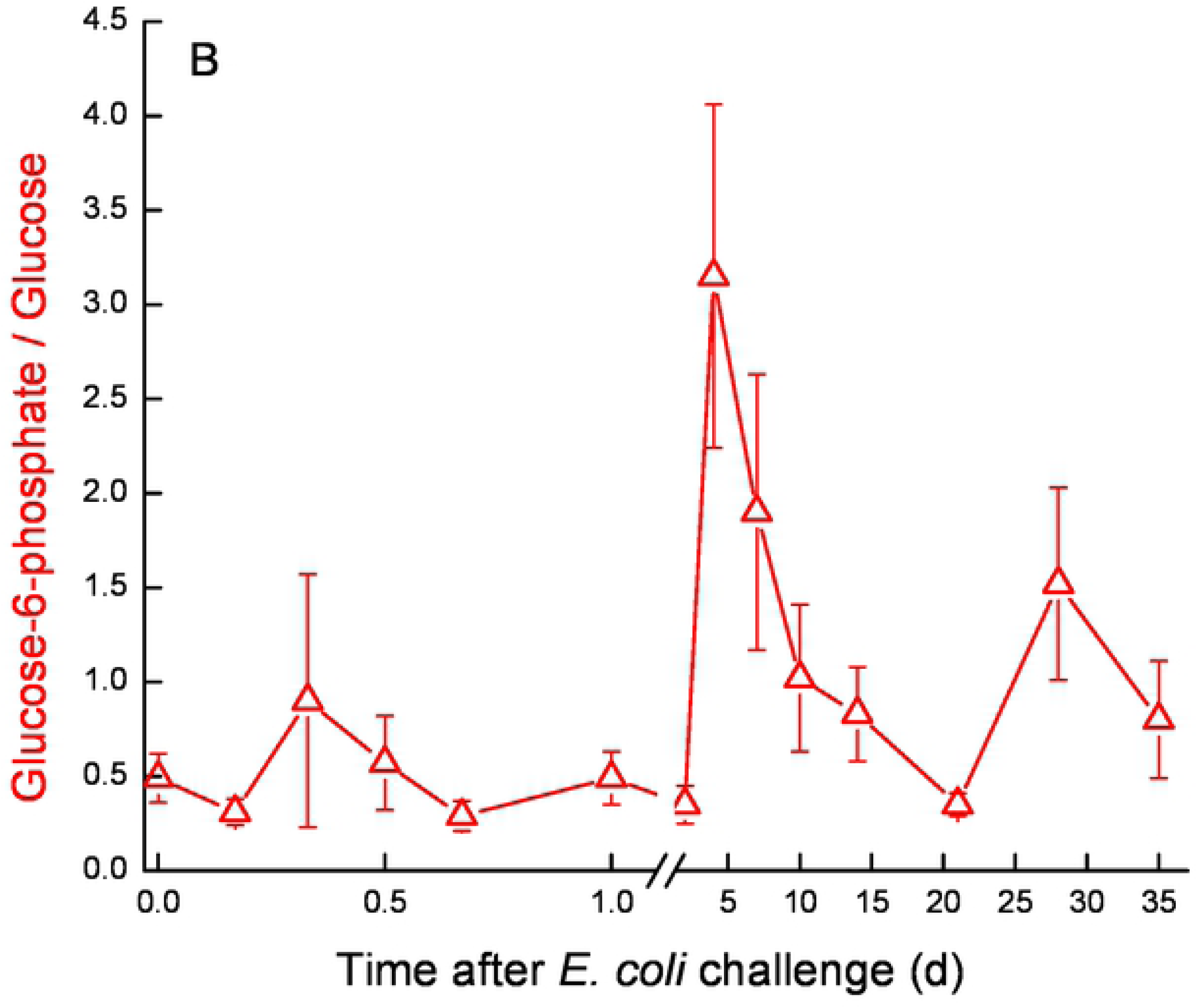
Mean and SE of glucose-6-phosphate (○; μM), glucose-6-hosphate/glucose (△) (Fig. 6A) and glucose-6-phosphatedehydrogenase (□; μM) (Fig. 6B) measured in the infected glands of 10 cows challenged with different mammary pathogenic strains of *E. coli*.

Oxaloacetate levels increased in a two-step way. A first significant increase was observed at ~16 h from ~0.8 mM to ~1.3 mM, then a second sharp increase was measured on 7 DPC to >1.75 mM after which oxaloacetate levels decreased again to ~1.3 mM and remained higher than pre-challenged levels (Fig. 7). In the infected glands, La and Ma concentrations displayed similar patterns. Their levels started to elevate at ~12 h, peaked on 2-4 DPC and declined thereafter (Fig. 8). A sharp increase of La was measured earlier, at ~12 h, whereas increase of Ma was detected at 1 DPC. La and Ma levels remained about 6 and 3 folds higher than pre-challenged levels until the end of the experiment, respectively. Pyruvate behaved differently, compared to the previous parameters. Pyruvate levels fluctuated during the first day post-challenge; gradually increasing until a peak > 1400 µM at 7 DPC, after which Pyr levels sharply declined to ~750 µM at 10 DPC. By the end of the study, Pyr levels were lower than pre-challenged levels (~600 µM and ~900 µM, respectively) (Fig. 9). Citrate concentration dynamics mirrored that of La and Ma above. Citrate levels sharply decreased at 16 h from 8.9 mM to lowest levels (3.8 mM) at 2 DPC, then gradually increased but remained lower on 35 DPC compared to pre-challenged levels (Fig. 10). Consequently, the ratio Cit/La + Ma dropped sharply, starting at 12 h from ~17 to < 1 at 2 DPC and then started to rise to the pre-challenged level ratio at 28 DPC (Fig. 11).

**Figure 7.**
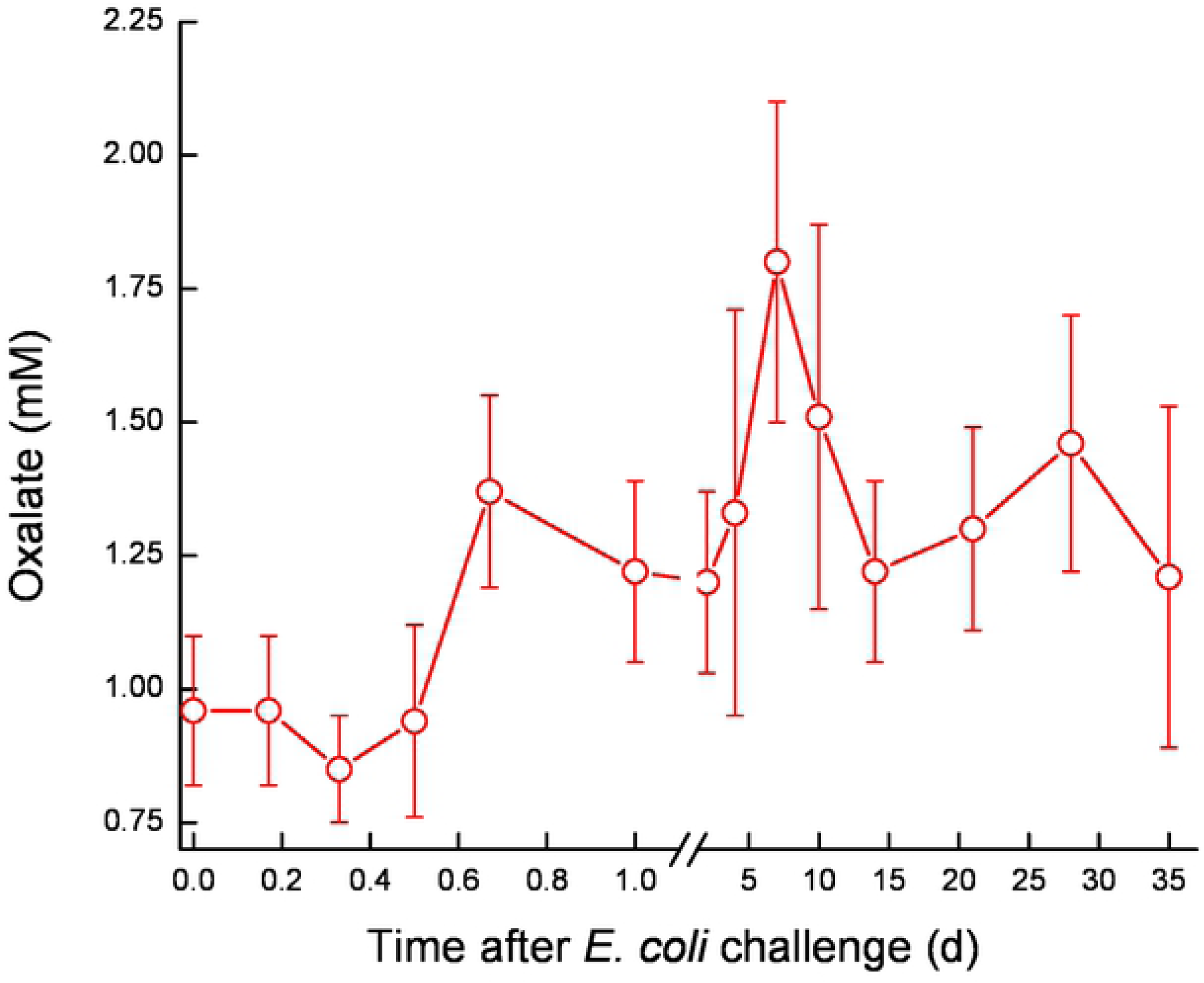
Mean and SE of oxalate (mM) measured in the infected glands of 10 cows challenged with different mammary pathogenic strains of *E. coli*.

**Figure 8.**
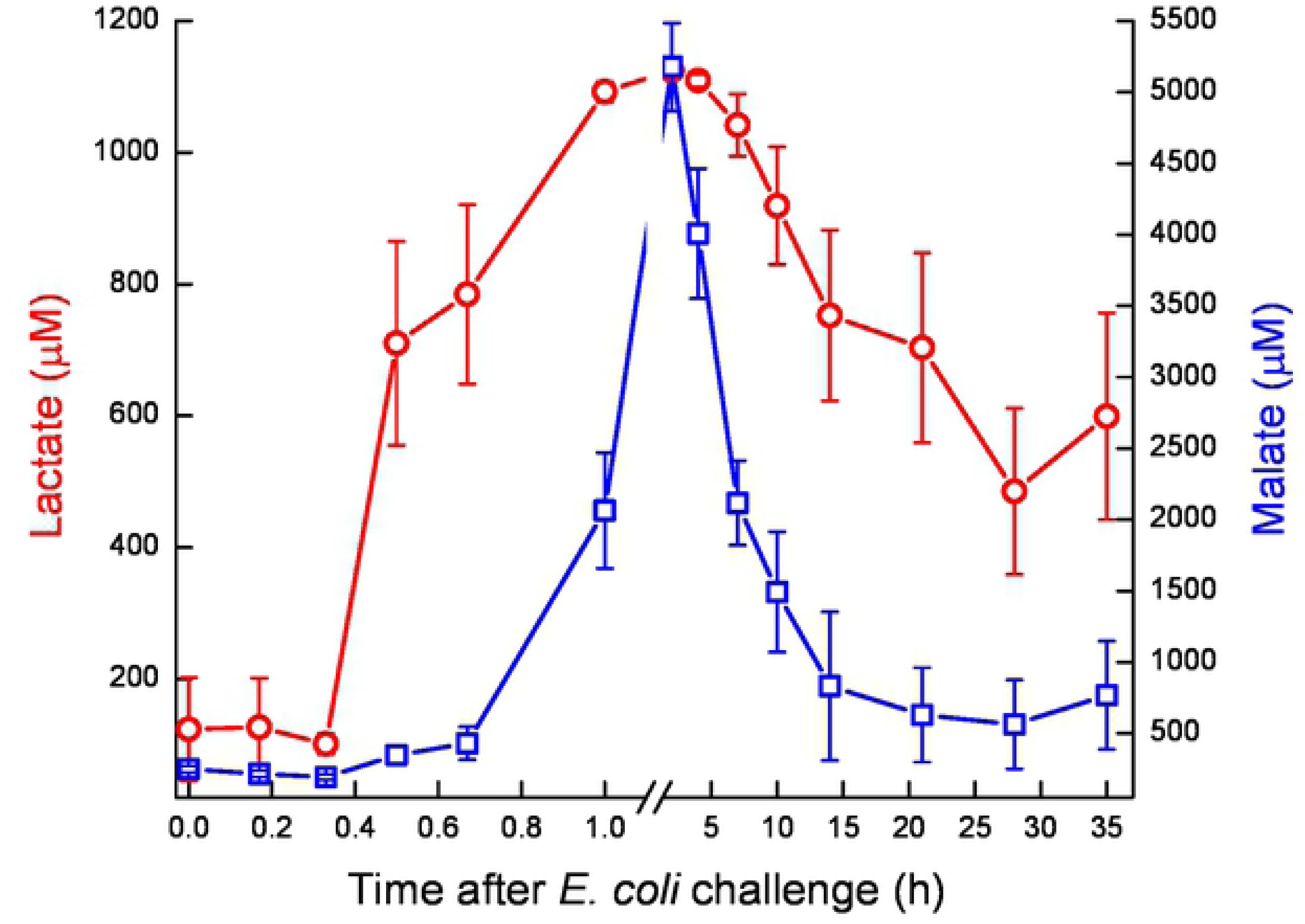
Mean and SE of lactate (○) and malate (□) measured in the infected glands of 10 cows challenged with different mammary pathogenic strains of *E. coli*.

**Figure 9.**
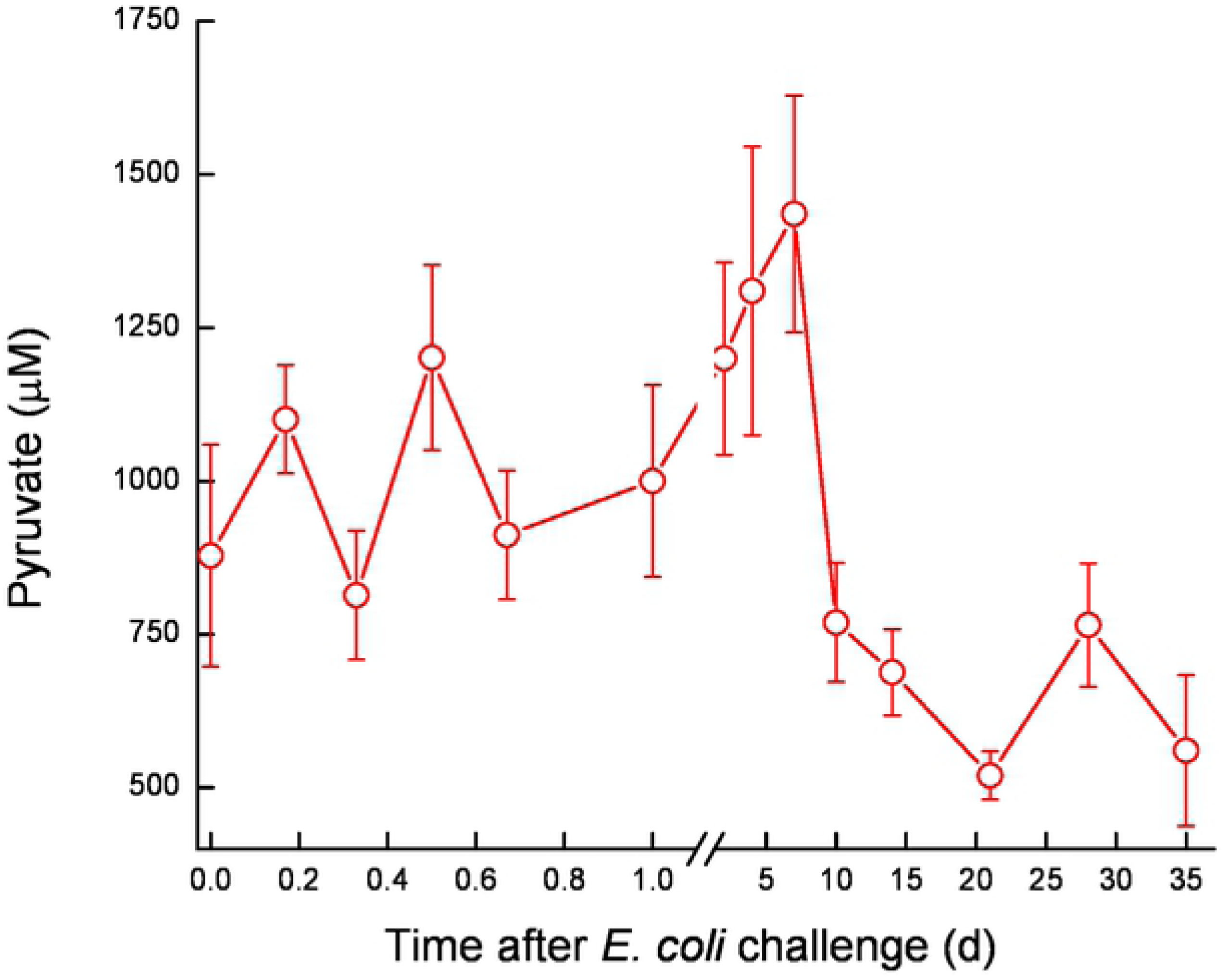
Mean and SE of pyruvate (μM) measured in the infected glands of 10 cows challenged with different mammary pathogenic strains of *E. coli*.

**Figure 10.**
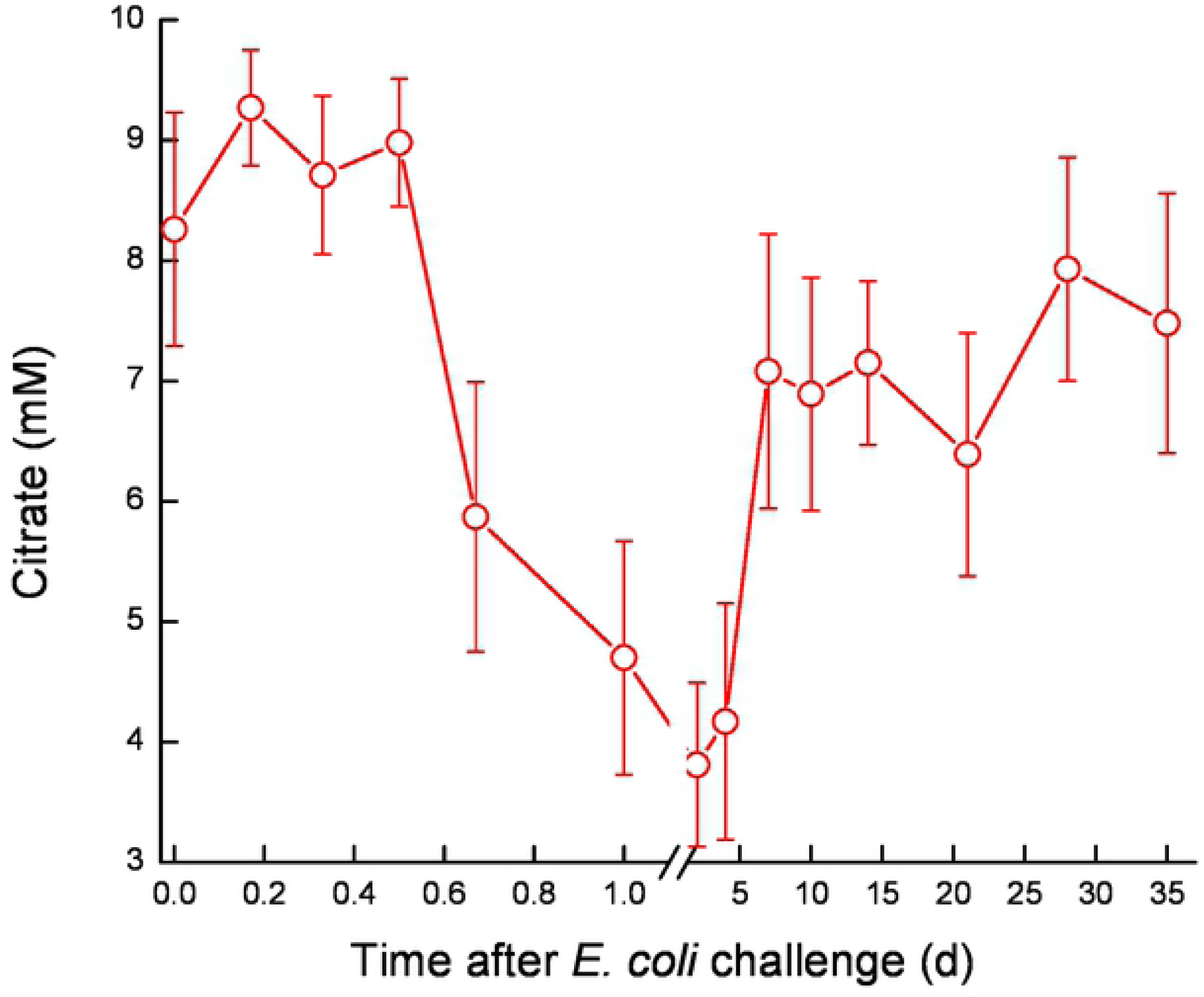
Mean and SE of citrate (mM) measured in the infected glands of 10 cows challenged with different mammary pathogenic strains of *E. coli*.

**Figure 11.**
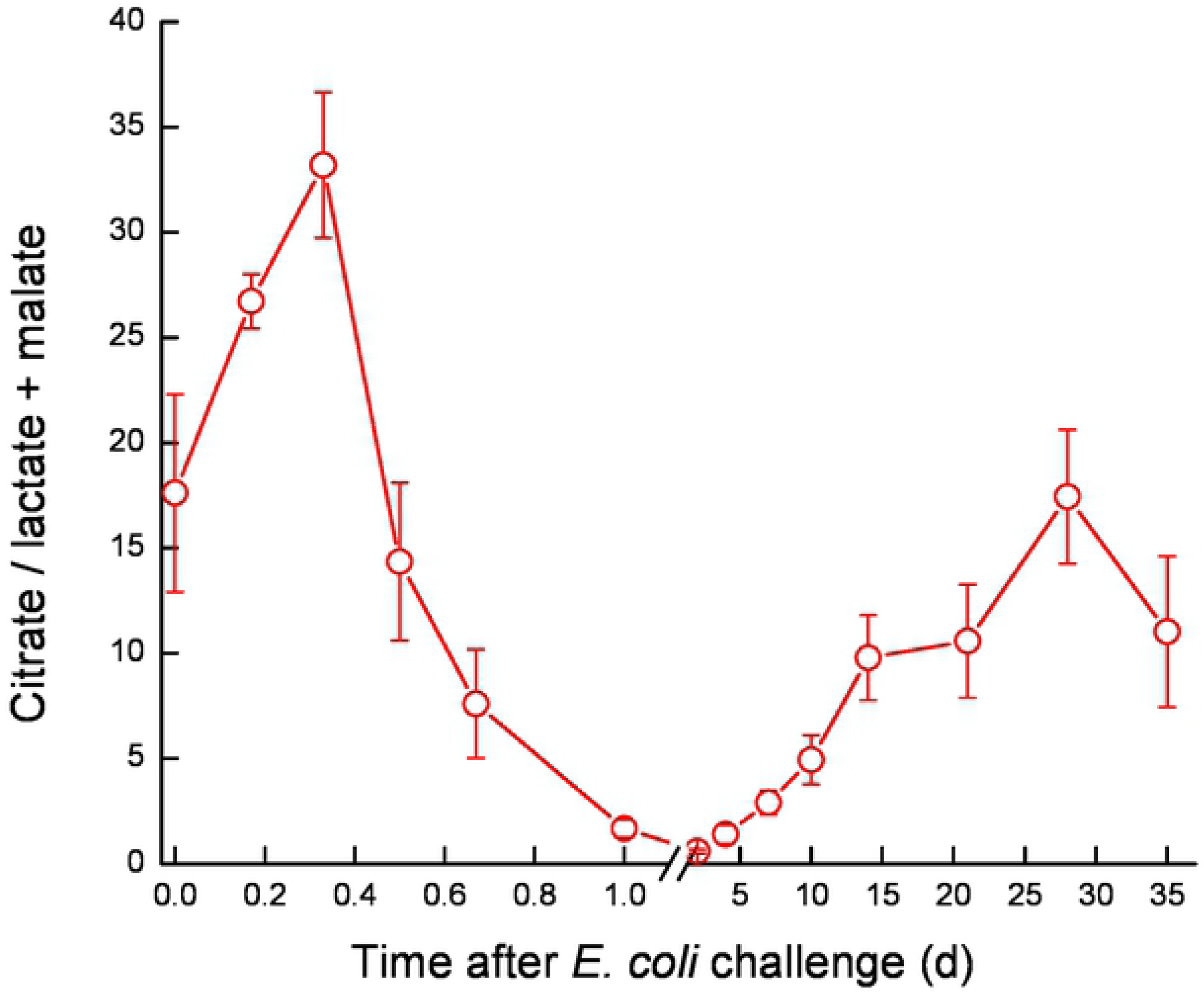
Mean and SE of the ratio citrate / lactate + malate measured in the infected glands of 10 cows challenged with different mammary pathogenic strains of *E. coli*.

Catalase and lactoperoxidase activity, LDH, BSA and nitrite concentrations displayed similar patterns. Their levels started to elevate at ~12 h PC, peaked at 2-4 DPC and declined to about pre-challenged levels at the end of the study (Fig. 12A-E). These blood proteins, enzymes and ion significantly positively correlated with SCC and milk protein, and significantly negatively correlated with infected glands and cow milk yield, lactose, % casein and CF (S1 Fig.). Lactoferrin concentration increased in 16-24 h from ~300 to 1,600 µg/mL, peaked at 7 DPC and declined to the end of the study, but remained ~ 2 fold higher from the pre-challenge level (Fig. 12F). IgG concentration increased within 12 h from 0.2 to >2 mg/mL, peaked at 72 DPC > 7, remained in this level up to 21 DPC and only then declined, but also remained ~2 fold higher from the pre-challenged level at 35 DPC (Fig. 12G).

**Figure 12 A-G.**
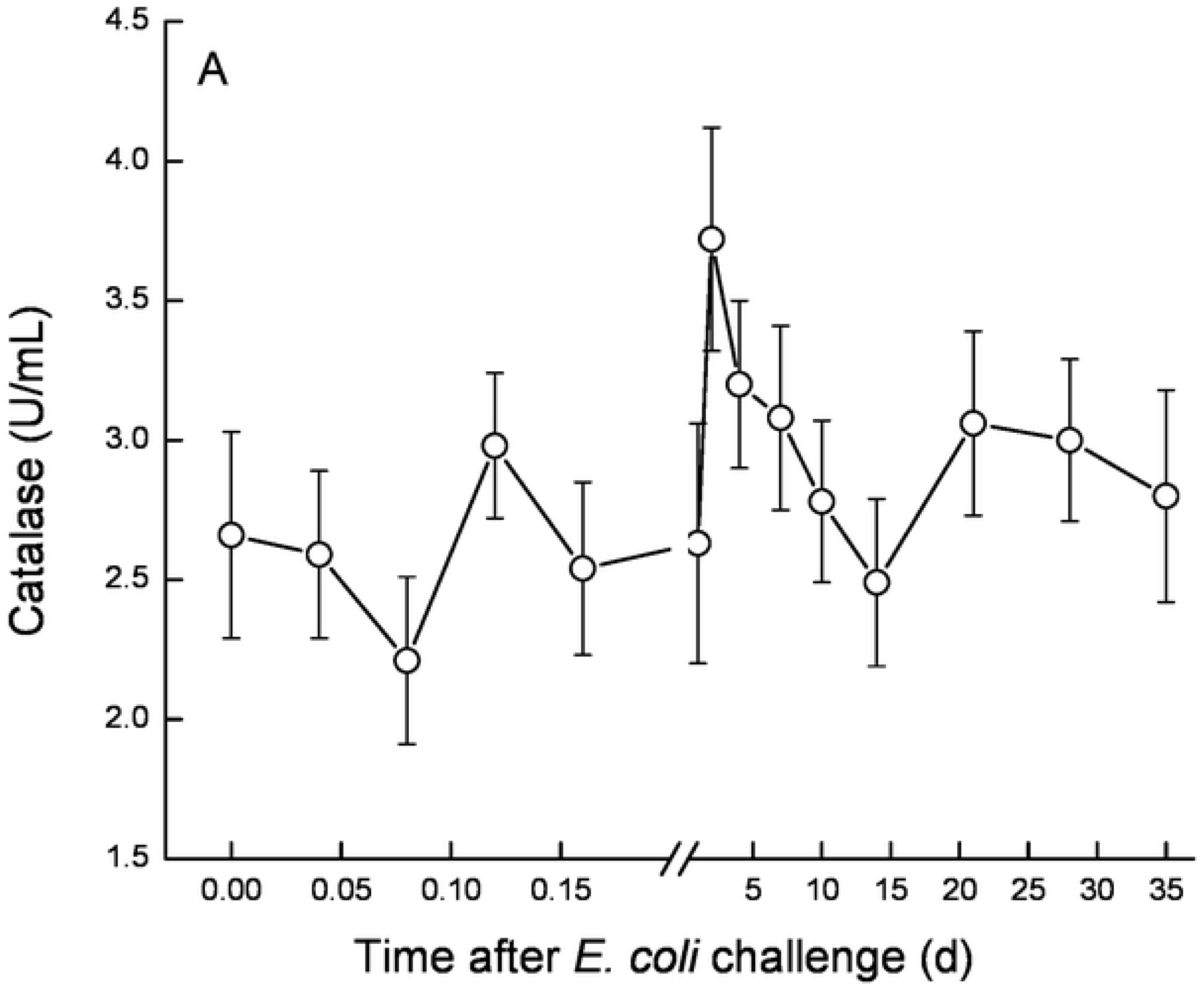
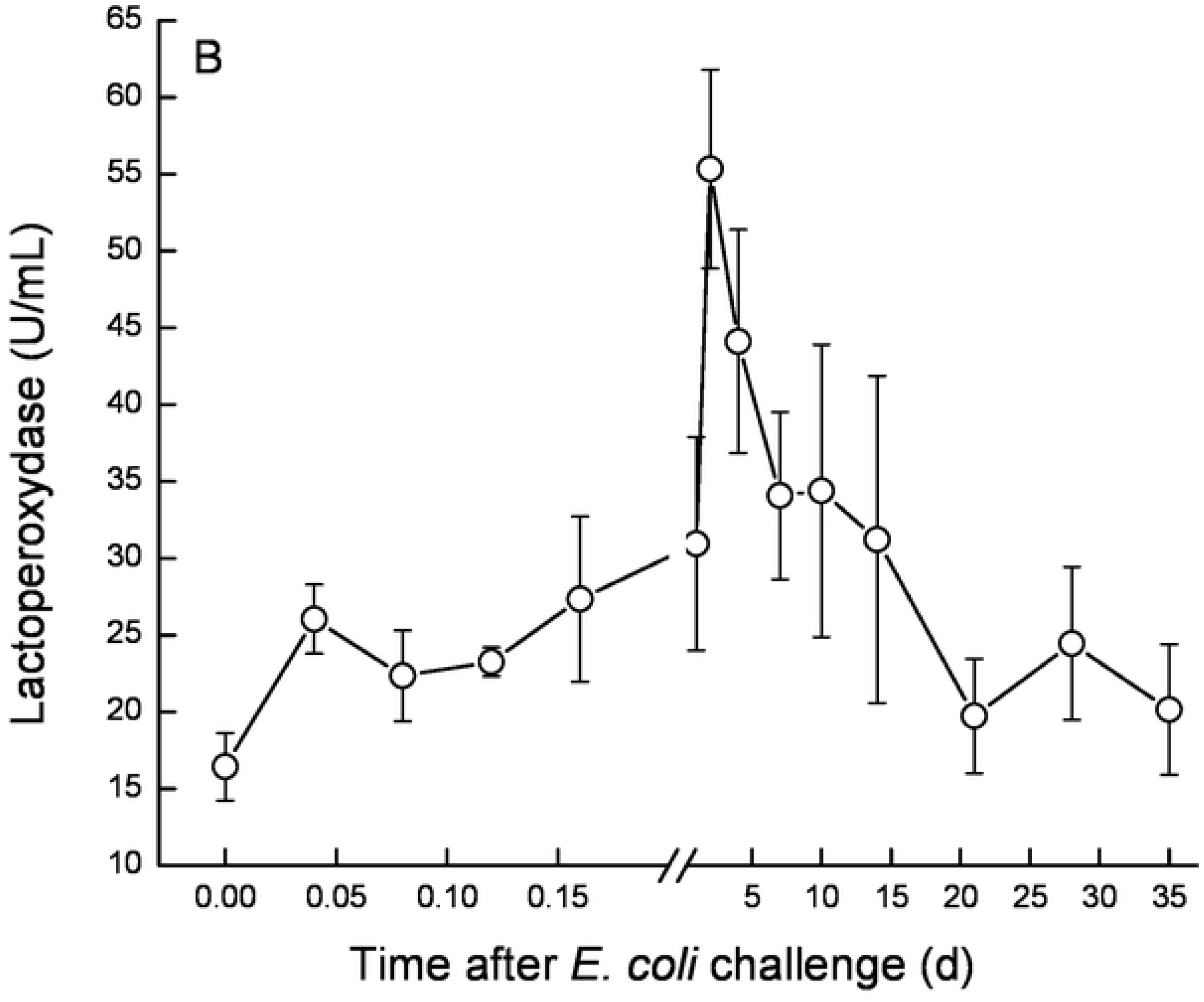
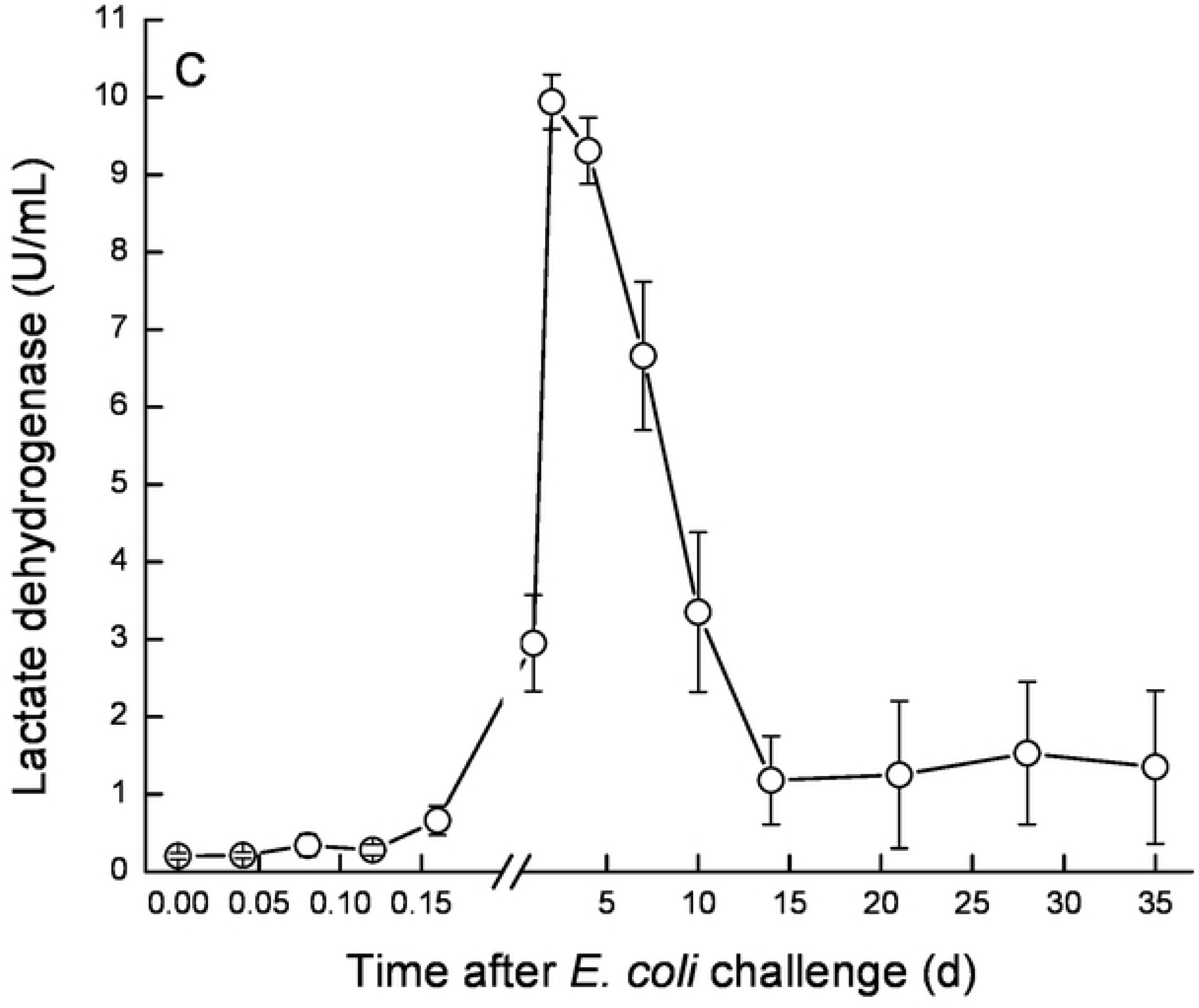
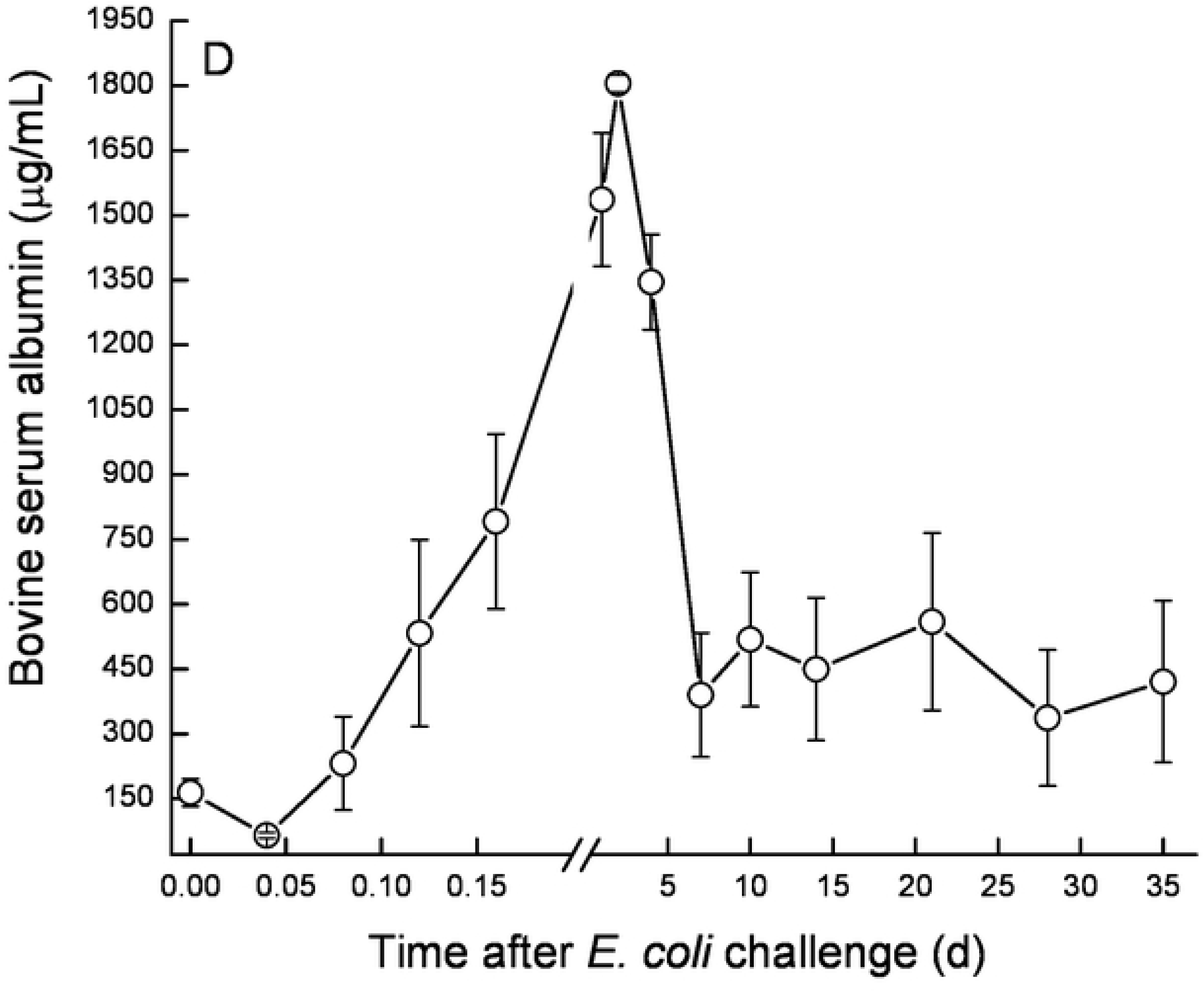
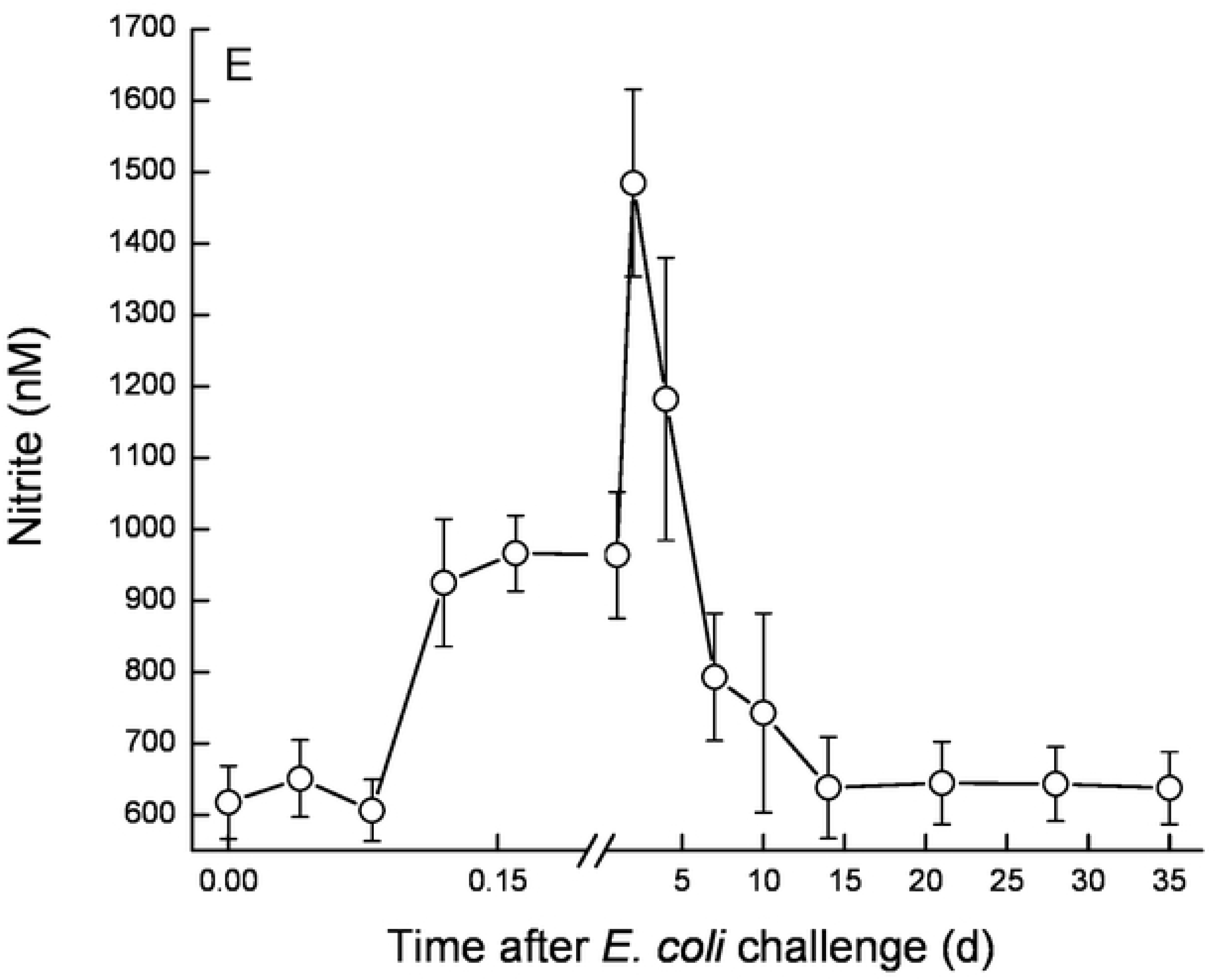
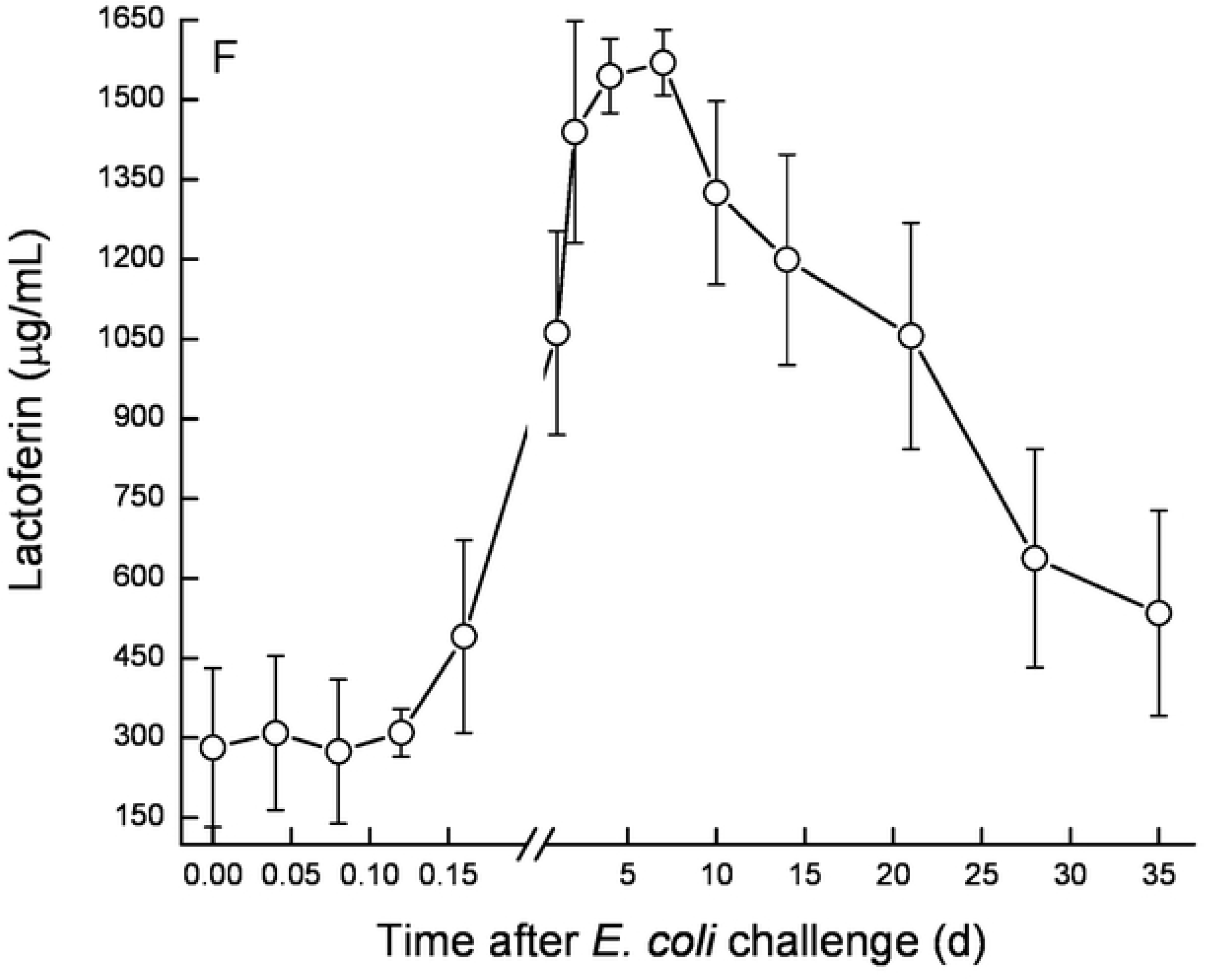
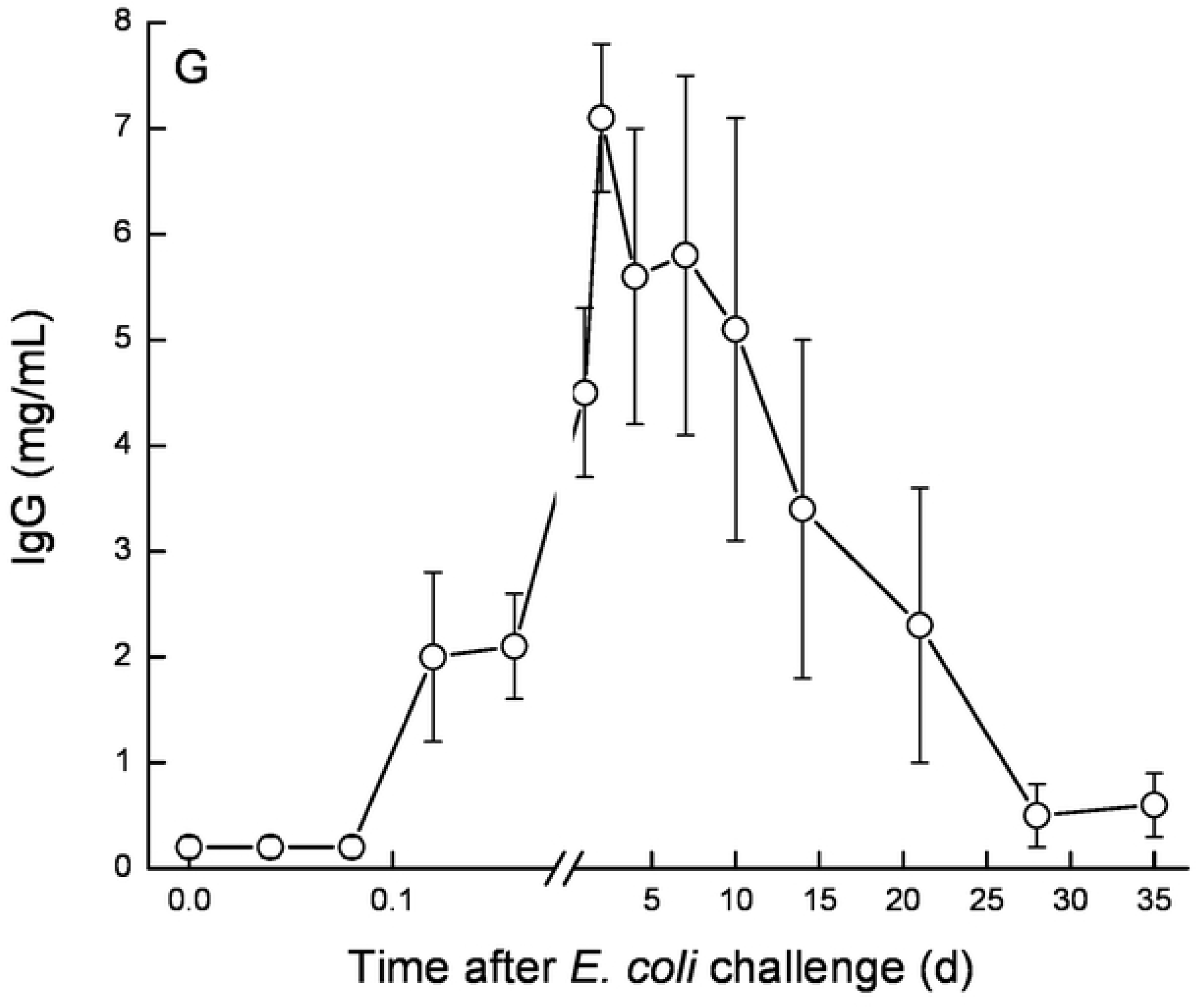
Mean and SE of IgG (mg/mL) (Fig. 12A), Nitrite (nM) (Fig. 12B), bovine serum albumin (µg/mL) (Fig. 12C), lactoferrin (µg/mL) (Fig. 12D), lactate dehydrogenase (U/mL) (Fig. 12E), lactoperoxidase (U/mL) (Fig. 12F) and catalase (U/mL) (Fig. 12G) in the infected glands of 10 cows challenged with different mammary pathogenic strains of *E. coli*.

### Histology at 42 DPC

Histological changes in infected and control glands were compared within each and among cows. Overall, the major differences in the control glands was the level of alveolar cuboidal epithelial structure in full lactating glands in the high yielding cows (Fig. 13A), and increased interlobular collagen rich areas with fibrous stroma and fat in low yielding cows (Fig. 13B). In challenged glands, regardless if bacteria were isolated on 35 DPC or not, increased interlobular collagen rich areas with fibrous stroma and fat were observed, and in the high yielding cows, normal alveolar structure. Moreover, in most challenged glands, leucocytes infiltration was observed, mainly inter-lobular mononuclear infiltration, and in two of the 10 glands PMN were observed within normal alveoli (Fig. 13C).

**Figure 13 A-C.** Representative hematoxylin-eosin histological sections of control and infected glands of 10 cows challenged with different mammary pathogenic strains of *E. coli*. Note the level of alveolar cuboidal epithelial structure in full lactating glands in high yielding cows (Fig. 13A), and increased interlobular collagen rich fibrous stroma and fat in low yielding cows (Fig. 13B). In challenged glands, increased interlobular collagen rich areas with fibrous stroma and fat was observed, with mononuclear leucocytes among lobules and PMN within normal alveoli in two out of 10 glands (Fig. 13C).

## Discussion

*E. coli* mastitis is characterized by acute inflammation, including local clinical signs of inflammation, such as swelling and pain, and sharp decrease in milk production along significant changes in its composition, such as increased SCC and passage of blood elements into milk. Systemic signs, including increased body temperature, diarrhea and dehydration are sometimes also observed. *E. coli* growth in milk in the mammary gland is fast, and increased bacterial counts are detected as early as within 4 h from challenge [4] even when a minimal inoculum is used (10-30 CFU in the present study) to closely represent natural infections. The present study focused on the physiological and biochemical changes in milk following challenge with three different MPEC strains isolated from distinct mastitis presentations (VL2874, VL2732 and P4 [4]. Although some significant differences were found in the inflammatory and immune responses in the mammary gland between these strains, the physiological and biochemical revealed only minor differences without statistical significance. Therefore, all three strains were combined during analysis.

Cows did not receive medication, thus bacteria clearing in milk was natural, starting on 7 DPC. In 50% of the glands, bacteria shedding was intermittent until the end of the study. Regardless of the clearing rate and bacteria numbers in milk, no significant correlation was found with all the parameters tested, suggesting that the recovery process depends more on the severity of the infection and the inflammatory response during the first hours or days after *E. coli* entry in the mammary gland.

Two clear infection stages were observed: 1: Initiation and establishment of the infection, which started at 4-10 h PC and ended by 7-15 DPC; 2: Resolution of infection, as expressed by most of the parameters tested recovering to pre-challenge levels, although with different dynamics. However, of note, SCC and milk yield at the infected glands and at cow level was not restored until the end of the study at 35 DPC, similar to our observations in natural *E. coli* mastitis [5].

Milk production in the mammary epithelial cells exists under osmotic balance between blood and the epithelial cells [20]. Acute signaling of *E. coli* LPS and immune cytokines like IL8, TNF and others occurs within hours, starting a cascade of events, including infiltration of leucocytes, mainly PMN, and selectively opening of the tight junctions [21] and has been showed in our earlier study [4].

The shift of mammary epithelial cells metabolism to anaerobic glycolysis as a tradeoff between use of Glu to support lactose synthesis and liberation of Glu to support the immune system was thorough fully discussed by Silanikove et al. [6] using *E. coli* LPS as a model for mammary gland infection. LPS is a major immunogenic determinant of *E. coli* and Gram-negative bacteria in general, but there are important differences between LPS inoculation vs. and live *E. coli* bacteria challenge. First, a single inoculation of LPS leads to a single triggering event for the inflammatory response, whereas live bacteria replicate in the mammary gland and therefore represent a longer trigger for inflammation. Second, LPS by itself does not cause direct damage to the gland tissues whereas live bacteria have different virulence factors that can interact and damage the mammary epithelial cells, leading to longer recovery process. Third, the amount of LPS often used to induce mastitis may be higher than the amount of LPS found in small bacterial inoculum, such as the one used here. Here we observed that the inflammation caused by the *E. coli* could be divided into two stages: an acute phase, with clinical symptoms, and a chronic phase, independent of bacteria clearance, and in response to the tissue damage caused in the first phase (Fig. 14). The present results suggest that the tight junctions in the mammary alveoli were “open” at 12-24 h PC. This is expressed by infiltration of Glu, G, IgG, BSA, Nitrite, LDH from blood into the milk for 2-3 d after LPS inoculation [22] and 6-10 d after live bacteria inoculation. However, the free transfer of the immune cells (SCC) remained for weeks, suggesting controlled infiltration. Overall the dynamics of biochemical parameters evaluated was similar to that of LPS induced mastitis [6], but levels achieved were either higher (e.g. glucose-6-phosphate and lactate dehydrogenase activity) or changes remained longer (e.g. lactate dehydrogenase, Glu6P/Glu,). The ratio Cit/La+Ma, which represents the milk-reflected mitochondrial/cytosol metabolic index (Silanikove et al., 2014), significantly decreased during the acute phase of inflammation, as described in LPS induces mastitis. Here, however, this change remained for a longer period, and the ratio did not return to pre-challenge levels by the end of the study at 35 DPC. Cows tested 30 days after an episode of natural *E. coli* mastitis (“post-*E. coli*”) showed similar results (Silanikove et al., 2014). Peak bacterial counts in milk were observed at 24 h [4], following a steep decrease in citrate between 16 to 24 h after challenge and concomitant to a significant increase in lactoferrin in milk. The ability to assimilate iron from citrate is a key feature of MPEC [3] and therefore it is possible that the exchange of citrate to lactoferrin as main iron chelator in milk contributes to reduce bacterial load.

Interestingly, regardless of bacteria presence in the challenged mammary glands, milk yield was lower than pre-challenge, the number of the immune cells remained high (>1×10^6^ cell/mL) and the quality of the milk was low as expressed by impaired clotting parameters, even though fat and protein levels were actually higher than pre-challenged ones. This could be related to other milk constituents, but also to the quality of fat and protein fractions present in milk following inflammation, which deserve further investigation. Histology at 42 DPC allowed the observation of disrupted mammary gland tissues, affected alveolar cuboidal epithelial structure and increased interlobular collagen rich areas with fibrous stroma and fat. In most gland, leukocytes infiltration, mainly mononuclears among lobules and PMN within the normal alveoli was present. This correlates with the chronic, long-term effects reminiscent of mastitis caused by *E. coli*, indicating tissue damage and the process of cleaning of damaged tissue, and which result in affected milk quality and yield [5]. Overall, the present study adds information about the physiological and biochemical changes observed in milk during and following acute intra-mammary inflammation induced by infection by *E. coli* bacteria. Moreover, the present results highlight the shift between the acute phase and a chronic phase of *E. coli* mastitis, the latter being independent of bacterial shedding in milk; and is therefore independent from an ongoing infection. This chronic phase of inflammation is characterized by continuous changes in milk properties for a long period after clinical recovery, and the present results may explain the long-term effects on milk quality that have been previously described [5]. This study also emphasizes some differences between LPS and live bacteria-induced mastitis, which are important to be accounted when choosing the right model for mastitis research.

## Conclusions

*Escherichia coli* challenge induces acute conversion of epithelial cells metabolism from principally mitochondrial-oxidative to principally cytosolic (glycolytic), which allows diversion of metabolic resources normally used to synthesize milk and to support the immune system. In turn, reduction in lactose concentration is consistent with the concept that it is part of the defense mechanism of the mammary gland. We suggest a two-phase mechanism for the development of *E. coli* mastitis. In phase 1, the acute phase, bacterial infection in the mammary gland or in LPS induced mastitis, triggers an immediate immune and inflammatory responses in which the endothelial-mammary barriers are open, allowing for cellular and humoral infiltration from the blood to the milk. In phase 2, the chronic phase, tight junctions close and infiltration is limited to areas of deep damage in the gland, which may perpetuate alterations which can be measured in the milk, regardless of bacterial clearance from the mammary gland.

